# Phylogenomics of novel clones of *Aeromonas veronii* recovered from a freshwater lake reveals unique biosynthetic gene clusters

**DOI:** 10.1101/2024.05.08.593017

**Authors:** Opeyemi U. Lawal, Noah Bryan, Valeria R. Parreira, Rebecca Anderson, Yanhong Chen, Melinda Precious, Lawrence Goodridge

**Affiliations:** Canadian Research Institute for Food Safety (CRIFS), University of Guelph, Guelph, ON, Canada N1G 2W1; Bayview Secondary School, 10077 Bayview Ave, Richmond Hill, ON L4C 2L4

**Keywords:** *Aeromonas*, genomics, antimicrobial resistance, biosynthetic gene cluster, water quality, freshwater lake, public health

## Abstract

Aquatic ecosystems are important reservoirs for clinically relevant pathogens and antimicrobial resistance genes, thus present a significant risk to global health. Here, we assessed the phylogenomics of *Aeromonas veronii* (*A. veronii*) recovered from Lake Wilcox in Ontario using a combination of morphological, biochemical, and whole-genome sequencing (WGS) techniques.

Eleven distinct bacterial colonies were isolated and identified as *A. veronii* (n=9), and two other *Aeromonas* species (*A. caviae* and *A. allosaccharophila*), with significant discrepancies noted between biochemical and WGS identification methods. Of note, 67% (n=6/9) of *A. veronii* isolates were human pathogens (Pathogenicity score ≥ 0.50). The genomic analysis revealed high genetic diversity among the *A. veronii* isolates, including the discovery of 41 novel alleles and seven new sequence types (ST) suggesting the lake as a reservoir for multiple human pathogenic clones of this bacterium. The comparison of the newly isolated and sequenced *A. veronii* with 214 *A. veronii* genomes revealed significant genetic diversity and suggests potential broad geographical dissemination of strains. Chromosomal genes (*OXA-912* and *cphA* [*cphA3, cphA4, cphA7*]) genes encoding resistance to β-lactamases were detected in all isolates. Human and non-human pathogenic strains of *A. veronii* differed in their virulence gene content, with type III secretion systems being associated with human pathogenic isolates. Mobilome analysis revealed the absence of plasmids in *A. veronii* isolates and the presence of 13 intact the great majority of which were P22-like (Peduoviridae) phages, and nine different insertion sequence families. Novel biosynthetic gene clusters were identified and characterized, indicating the potential for unique secondary metabolite production in *A. veronii* with different pathogenic potential. Overall, this study underscores the importance of continuous surveillance of aquatic ecosystems for the presence of pathogens, contributing to our understanding of their evolution, potential for human pathogenicity, and the ecological roles of their genetic elements.

## BACKGROUND

The role of aquatic ecosystems as reservoirs for clinically relevant pathogens and antimicrobial resistance genes (ARG) has recently gained attention as the importance of assessing the quality of these ecosystem is paramount in public health (1, 2). Freshwater bodies like lakes and rivers used for recreational purposes can significantly impact the health of communities (3, 4). Poor water quality in these settings poses a substantial risk for the transmission of various waterborne diseases, including pathogenic viruses, protozoa, and bacteria including *Aeromonas* species that thrive in such contaminated water (2–4).

*Aeromonas* species are Gram-negative, facultative anaerobic rods, found in various aquatic environments (5–7), and known for their ability to survive in diverse environments, ranging from freshwater to the intestinal tracts of animals (5, 7). While some *Aeromonas* species including *Aeromonas salmonicida*, *Aeromonas hydrophila*, and *Aeromonas veronii* are known fish pathogens, *A. veronii* is one of the four species that are considered as potential human pathogens (8–10). *A. veronii* is an emerging human pathogen causing a wide range of diseases in human and animals including gastroenteritis, respiratory and skin infections and septicemia (9–11). In addition, *A. veronii* is increasing being recognized as a significant concern to food safety due to its frequent presence in different types of food, particularly in minimally processed ready-to-eat seafood (12, 13). Of note, the frequent and global occurrence of highly virulent strains of *A. veronii* has been detected in food samples such as meat, milk, catfish and fish in countries including Brazil (13), Egypt (14), India (15), Israel (11), and the United States (16, 17), among others. The adaptability of *A. veronii* to various conditions poses a challenge for water quality management, especially in environments with high anthropogenic activities, where the bacterium can be a potential source of infection (5, 18).

The mechanisms of pathogenicity of *A. veronii* involve the production of various toxins and virulence factors that contribute to its ability to infect host cells and cause disease (5, 18). A significant concern with *A. veronii* is its capacity for antimicrobial resistance (7, 19, 20). The presence of antimicrobial-resistant strains in aquatic environments is a public health concern, as it not only affects the treatment of *Aeromonas*-related infections but also represents a potential reservoir for the spread of resistance genes to other pathogenic bacteria (7, 19, 20). Studies on the population structure of *A. veronii* have described genetic diversity driven by its adaptability to various environmental conditions. These factors could drive variability in strains regarding pathogenicity and resistance to environmental stresses in this bacterium, with practical implications for public health and water management (5, 8, 18).

In recent years, advancements in sequencing technologies have greatly enhanced the genomic surveillance of known and emerging pathogens, such as *A. veronii*, across different environmental matrices(5). Despite these technological advancements, little importance has been given to *A. veronii,* especially in terms of its presence in freshwater, its impact on water quality, and its role in the dissemination of antimicrobial resistance (AMR) in both the environment and the food chain. Understanding the genomic surveillance and population structure of this bacterium is crucial for developing effective infection treatment strategies and ensuring public health safety.

We have previously reported the detection of clinically relevant pathogens in Lake Wilcox, including novel strains of *Bacillus anthracis* (21), and *Vibrio cholerae* (22) isolated at different time points. In this study, we employed a combination of culture-based detection and whole-genome sequencing to assess the presence of *A. veronii*, its extensive genomic fingerprint, its population structure, and the genomic characterization of stress response genes in Lake Wilcox. The genetic relatedness of *A. veronii* isolates was assessed by comparing them with previously sequenced strains in public databases using a comparative genomic approach.

## METHODS

### Description of sampling site

Lake Wilcox is a small kettle lake located in Richmond Hill in Ontario (43°56’56.69” N, 79°26’9.45” W). Historically, the lake is used for recreational purposes by the surrounding community and tourists. Despite being impacted by feces of surrounding wildlife, recreational activities have continued, and users have reported skin rashes and gastrointestinal symptoms after recreational activities (https://projectboard.world/ysc/project/the-phage-takes-centre-stage-for-water-quality-testing) **(Figure 1)**.

**Figure 1.**
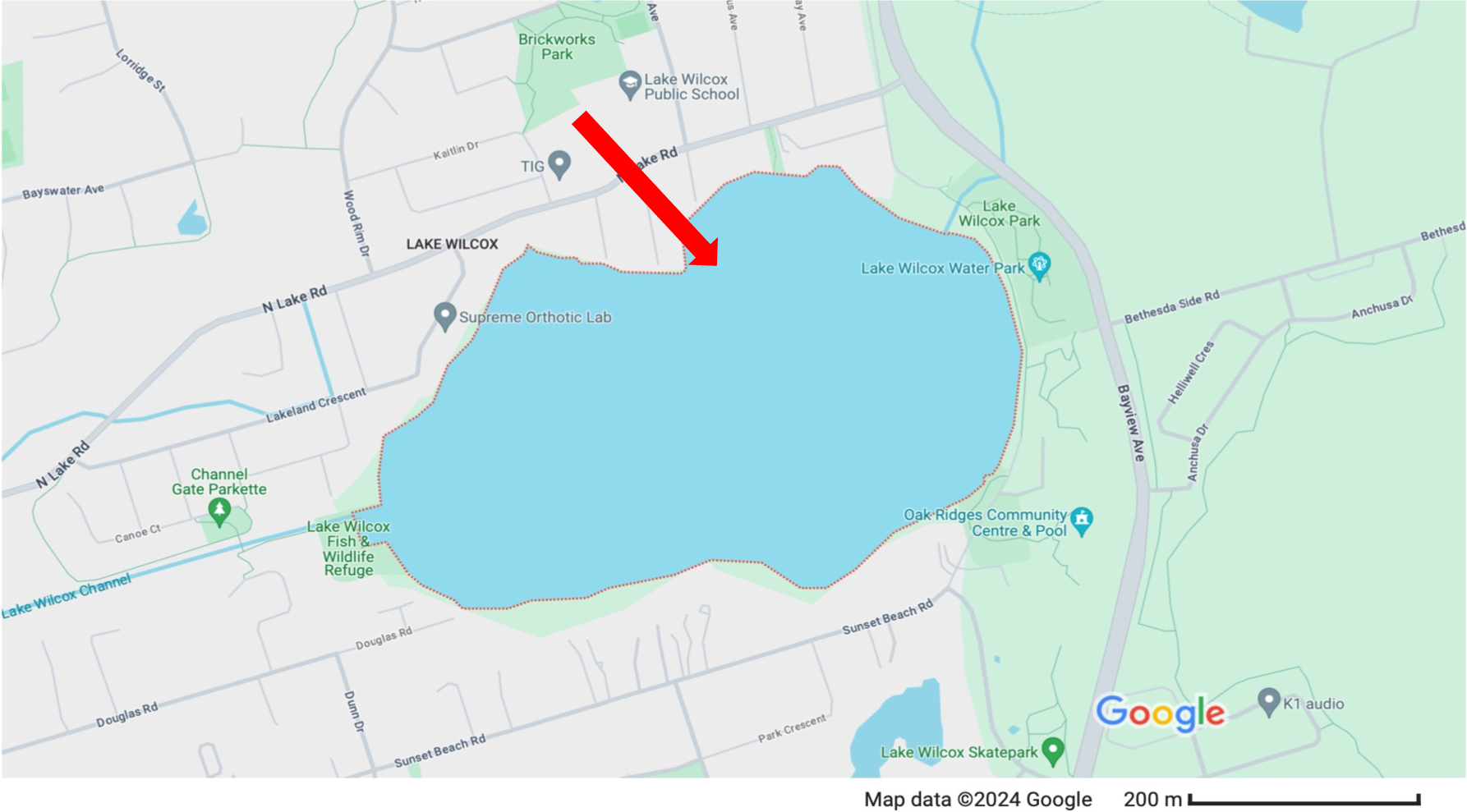
– Location of Lake Wilcox in Ontario, Canada. Image from http://maps.google.com

### Sample collection and processing

Freshwater samples were obtained from Lake Wilcox in the Summer of 2022 and Fall of 2023. Water samples were kept at 4° C and analyzed within 48 hours of collection. Samples were processed as described by Bryan et al, 2023 (23). Briefly, 1 mL of samples was serially diluted in 9 mL of lambda buffer (modified saline-magnesium buffer without gelatin) and plated onto tryptic soy agar (TSA). Following incubation for 24 h at 37° C, plates were analyzed for bacterial colonies. Distinct colonies of differing morphologies were sub-cultured onto TSA to obtain pure culture. The isolated colonies were *Gram* stained and taxonomic identification was performed using VITEK® (bioMérieux, Inc, Canada).

### Genomic DNA extraction and whole-genome sequencing

Genomic DNA from isolated colonies was extracted using the DNeasy blood and tissue kit (Qiagen Hilden, Germany) according to the manufacturer’s instructions. DNA libraries were prepared using the Illumina DNA prep tagmentation kit (#20018704) and IDT for Illumina DNA/RNA UD indexes (#20027213) following the manufacturer’s instructions. Paired-end (2 x 150 bp) sequencing was performed using the high output flow cell on the Illumina MiniSeq instrument as described previously (23, 24).

### Genome assembly and annotation

Raw paired-end reads were quality filtered using FastQC v0.11.9 (https://github.com/s-andrews/FastQC), and trimmed using Trimmomatic v0.39 (25). High quality reads with a Phred quality score above 20 were assembled *de novo* using the Skesa v2.4.0 pipeline (26). Assembly quality and genome completeness were assessed using QUAST v5.2 (27) and BUSCO (28), respectively. Genome annotation was performed using Prokka v1.14.6 (29).

### Gene content analysis

The antimicrobial resistance gene profile of all the isolates was determined using AMRFinder Plus v3.10.45 (30) and CARD (31) databases, while the virulence genes were identified using VFDB (32, 33). To define the mobile genetic elements of the collection, the draft genomes were screened for plasmids and prophages using MOB-suite v3.1.6 (34) and PHASTEST (35), respectively. Biosynthetic gene clusters were assessed using the antiSMASH v6 pipeline (36).

### Phylogenetic analysis

To construct the phylogeny, pangenome was generated from the annotated genomes using Roary v3.13.0 (37), core genome-based phylogenetic tree was constructed using FastTree (38). The general time reversible model was performed with 1000 bootstrap resampling for node support. Except as otherwise stated, all bioinformatics tools were executed using the default settings.

### Data availability

The whole-genome sequences reported in this study were deposited at DDBJ/ENA/GenBank under the BioProject accession numbers **<u>PRJNA893208</u>**. The raw sequence reads, and genome assembly accession numbers are listed in **Table 1**. In addition, accession numbers and associated metadata of genomes retrieved from NCBI are listed in **Table S3**.

## RESULTS

Bacterial species were isolated from the freshwater samples recovered from a freshwater lake over a period of one year using the spread agar plate method. Overall, 11 distinct colonies with different morphologies were selected and further characterized using morphological, biochemical-based, and whole-genome sequencing. Taxonomic identification using the VITEK® Compact system identified the colonies as *Aeromonas sobria* (n=8), *Aeromonas hydrophila/punctata* (n=2), and one isolate with an inconclusive result **(Table 1)**. Sequencing of the 11 isolates yielded 1,024,248 - 2,725,402 paired-ended reads per isolate **(Table 1)**. Using pubMLST and rMLST (39), as well as k-*mer*-based species taxonomic classification with the Kraken2 database (40), isolates were identified as *Aeromonas veronii* (n=9), *A. caviae* (n=1) and *A. allosacharophila* (n=1) **(Table 1)**. The average nucleotide identity (ANI) analysis with fastANI (41) showed that the nine *A. veronii* strains had >96% ANI when *A. veronii* GCF_000820225.1 strain was used as a reference, *A. caviae* strain NB-180 had 97.9% ANI with *A. caviae* GCF_000819785.1, while *A. allosaccharophila* had 96.21% ANI with the reference strain *A. allosaccharophila* GCF_000819685.1. The draft genomes of *Aeromonas* species yielded between 28 and 113 contigs, with a G+C content of 58 - 59 %, except for *A. caviae* that had a higher G+C content of 61.26%, a value that was comparable to the reference strain *A. allosaccharophila* GCF_000819685.1. The genome size was comparable between the three *Aeromonas* species identified and ranged between 4,390,436 and 4,690,056 bp, with >50× genome coverage **(Table 1**).

### Prediction of human pathogenicity of *A. veronii* sequenced

Considering that Aeromonas species are commonly associated with diseases in fish, we evaluated the potential of these isolates to be pathogenic to humans. We did this by comparing the proteins of the new strains with a database composed of protein families associated with either pathogenic or non-pathogenic organisms in humans, using the PathogenFinder tool. (42). Six out of the nine *A. veronii* isolates in this study had a pathogenicity score greater than 0.5, suggesting that they may be pathogenic to humans. Other isolates, including *A. caviae* and *A. allosaccharophila*, were predicted to be non-human pathogens **(Table 1)**.

### Population structure of *A. veronii* isolated from freshwater

To assess the genetic relatedness among isolates sequenced in this study, a combination of conventional MLST and whole genome-based phylogeny was employed. The *Aeromonas* MLST schema was used to determine the sequence types (STs) of all isolates. Of note, 41 novel alleles were identified among the 11 *Aeromonas* isolates and yielded nine unique allele profiles that were submitted together with the allele sequences and assigned to nine new STs (ST2530 – ST2538) **(Table 1 and Table S1)**. Two STs (ST2530 and ST2535) contained two isolates each while others were singletons suggesting the uniqueness of the isolates understudy and high genetic diversity in the population. The core genome SNP-based phylogeny of the nine *A. veronii* sequenced was constructed to using the complete closed genome of *A. veronii* AP022281.1 as reference. *A. caviae* and *A. allosaccharophila* were used as outgroups to root the tree. Isolates were grouped into two main clusters irrespective of the period of isolation **(Figure 2)**. Isolates were distantly related by SNPs with ≥100 SNPs difference **(Table S2)** except for a pair of isolates from different timepoints (NB-2/NB-4, Summer, 2022; and NB-178/NB-181, Fall, 2023) that were highly related differing only by 9 and 11 SNPs, respectively **(Figure 2)**. Of note, the SNP-based clustering observed was similar to the MLST-based population structure suggesting a good concordance between these methods for typing *A. veronii.* Overall, the high genetic diversity observed in this study suggest that the freshwater lake could serve as a reservoir for multiple strains of *A. veronii* that are pathogenic to humans.

**Figure 2.**
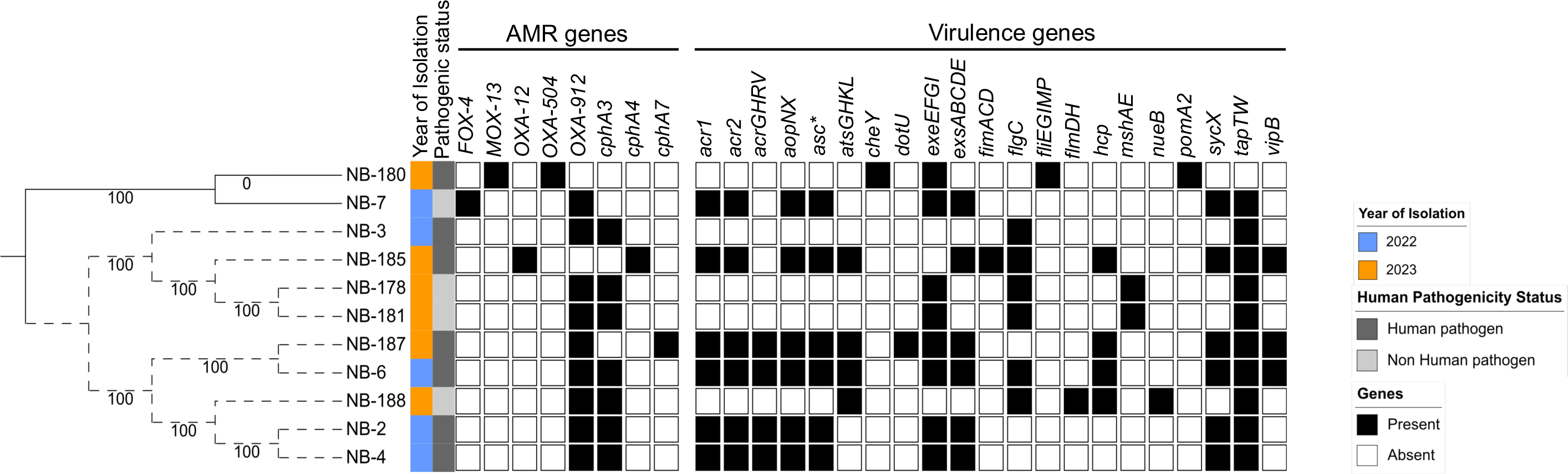
– Maximum likelihood tree of *Aeromonas veronii* recovered from freshwater lake in Ontario. Core genome sequence alignment was generated from draft genomes using MAFFT v7.453, and the tree was constructed using FastTree v2.1.11 using generalized time reversible nucleotide substitution model with 1000 bootstraps for node support. The dotted line depicts *Aeromonas* veronii. NB-7 (*A. allosaccharophila*) and NB-180 (*A. caviae*) were outgroups used to re-root the tree. The draft genomes were screened for genes encoding antimicrobial resistance and virulence using CARD and VFDB databases, respectively. The scale bar at the bottom represents nucleotide substitution per site. The tree was visualized using iTOL; (https://itol.embl.de).

### Global population structure of *A. veronii*

To assess the genetic relatedness of the sequenced isolates with global *A. veronii*, genomes and the associated metadata of 214 *A. veronii* in the *RefSeq* database (accessed on October 15, 2022) were downloaded and re-annotated (*see Method*). The 214 genomes were recovered from 18 different countries located in six continents between 1988 to 2022 from seven different sources including human, animal, aquatic ecosystem, fish, food, plant, and insect **(Table S3).** The pangenome size of the 214 *A. veronii* genomes together with the sequenced isolates (n=9) yielded 49,483 genes. A total of 2,248 core genes, defined as genes present in ≥95% of the genomes in the collection, were identified, whereas the shell and cloud genes totalled 2,294, and 44,941, respectively. The core genome-based maximum likelihood tree based on 2,248 core genes was constructed using *A. allosaccharophila* as outgroup to root the phylogenetic tree. The sequenced *A. veronii* isolates compared with global *A. veronii* species showed high genetic diversity which facilitated the clustering of the isolates into distinct clades **(Figure 3)**.

**Figure 3.**
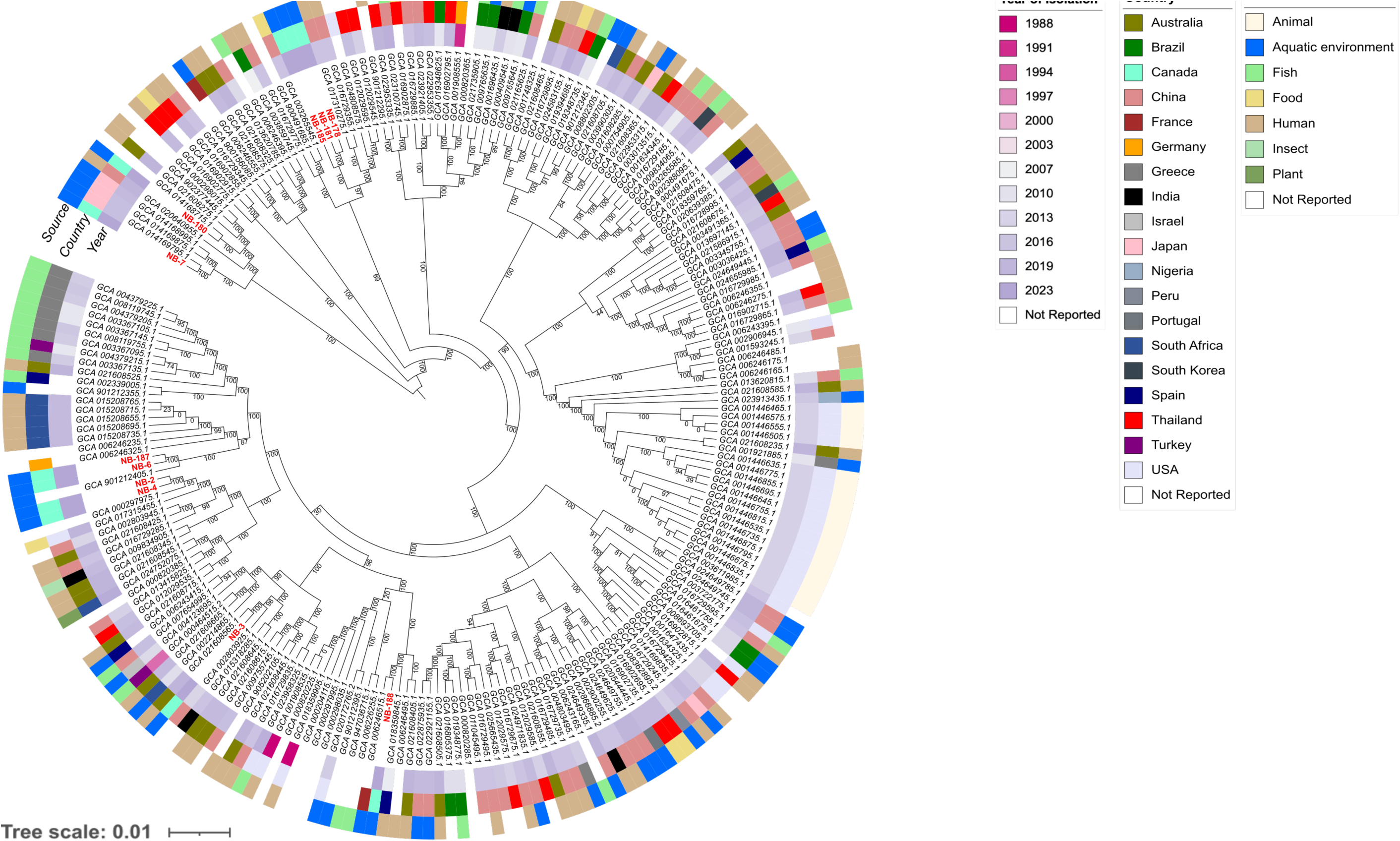
– Core-genome based maximum-likelihood tree of global *Aeromonas veronii* from different sources. Each node represents a strain. Core-genome sequence alignment was generated from draft genomes using MAFFT v7.453. Maximum likelihood tree was constructed using FastTree v2.1.11. The generalized time reversible nucleotide substitution was performed with 1000 bootstraps random resampling for support. *A. allosaccharophila* (NB-7) and *A. caviae* (NB-180) were outgroups used for re-rooting the tree. The scale bar at the bottom represents nucleotide substitution per site. The figure was generated using iTOL (https://itol.embl.de). *Aeromonas* isolates recovered in this study were labelled in red.

Isolates sequenced in this study were clustered into distinct subclades suggesting that they are distantly related to other global isolates. However, isolates from Turkey and Greece recovered in fish from different time points (2009, 2015 and 2016) were clustered together, a phenomenon that could suggest dissemination of *A. veronii* strains. Strain NB-188 belonged to the same subcluster as an isolate recovered from a similar aquatic system in France. Global *A. veronii* are distantly related. While some subclades were source-based, we could still observe a mixture of isolates from different countries and sources within others. Overall, *A. veronii* from different environments may have genetic signatures unique to pathogenic strains of this bacterium. This could also be important to determine or predict the source of isolates found in any matrix. Future evolutionary studies on this bacterium would be important to decipher this hypothesis.

### Stress response genes among the *A. veronii* sequenced

Genes encoding resistance to antimicrobials and virulence were assessed in the sequenced isolates. Genes encoding resistance to ý-lactams were detected in all the isolates. Different alleles of *cphA* (*cphA3, cphA4,* and *cphA7*) gene which belonged to the subclass B2 metallo-beta-lactamase that encodes resistance to carbapenem antibiotics were detected in all *A. veronii* isolates and in *A. allosaccharophila. OXA-912* that encodes resistance to penams, cephalosporins and carbapenems, and *cphA3* genes were predominant in the collection. These data suggest that *A. veronii* recovered from freshwater sources are a reservoir for β-lactamase resistance genes. Additionally, detection of virulence genes using a gene homology approach and a curated virulence gene database [VFDB (32, 33)] detected eight to 48 virulence genes in each sequenced *A. veronii* isolate. Relative to isolates predicted as human pathogens that contained 41 – 48 virulence genes (except NB-3), all the *A. veronii* predicted as non-human pathogens carried less virulence genes (≤9 virulence genes). Human and non-human pathogenic strains of *A. veronii* differed in term of their virulence gene content. While the flagellar and type IV pili associated genes involved in biofilm formation (18, 33) were detected in all isolates, type III secretion system (T3SS) associated genes were detected only in the isolates predicted as human pathogens **(Figure 2)**.

### Characterization of the mobile genetic elements among the sequenced *A. veronii*

Plasmids were not detected in any of the *A. veronii* isolates studied. However, *A. caviae* carried a small plasmid (3,976 bp, contig 44) that contains plasmid replication genes and uncharacterized proteins. Comparative sequence analysis with blast showed that the plasmid had the closest nucleotide sequence similarity (>99% coverage and identity) to *Aeromonas enteropelogenes* 9789_1_48 plasmid (<u>LT635650.1</u>). The detection and characterization of phage regions in the sequenced genomes yielded 13 unique intact phages among which four were predicted to be virulent phages (43). The completeness of all intact phage sequences was determined to be between 50 – 100% by CheckV (44). The phages were classified by PhaGCN (45) as Peduoviridae (n=10), and Chaseviridae (n=1) and two others unidentified according to the International Committee on Taxonomy of Viruses (ICTV) classification (46). In addition to *A. veronii* being predicted as host of the phages, other species of *Aeromonas* (*A. australiensis*, *A. diversa*, *Aeromonas* sp.) and *Serratia marcescens* could also serve as their hosts as determined by PhaBox (47, 48), suggesting that these phages could infect multiple hosts **(Table 2)**. Of note, the two pairs of isolates (NB-2/NB-4 and NB-178/NB-181) that were highly genetically related by SNP had the same phage content. No antibiotic resistance, toxin or related genes were detected in the intact phages. All the intact phages detected were screened for tailspike proteins (TSP) using TSPDB that contains 8,077 TSPs (49), but none was found. In addition to phages, other MGE identified in the collection include 16 different types of insertion sequence (IS) elements belonging to nine IS families **(Figure 4)**. IS*As19* (IS*481*), ISAs4 (IS*5*) were predominant in the sequenced *A. veronii.* The IS*4* subtype containing IS*Aeme13*, IS*As30*, IS*Apu1* was detected in isolates recovered from early sampling dates (Summer, 2022), whereas IS subtypes IS*630* (IS*Ahy2*), IS*200*/IS*605* (IS*As26*) and IS*21* (IS*As29*) were observed in isolates recovered in the Fall of 2023, suggesting a temporal distribution of IS elements in *A. veronii*.

**Figure 4.**
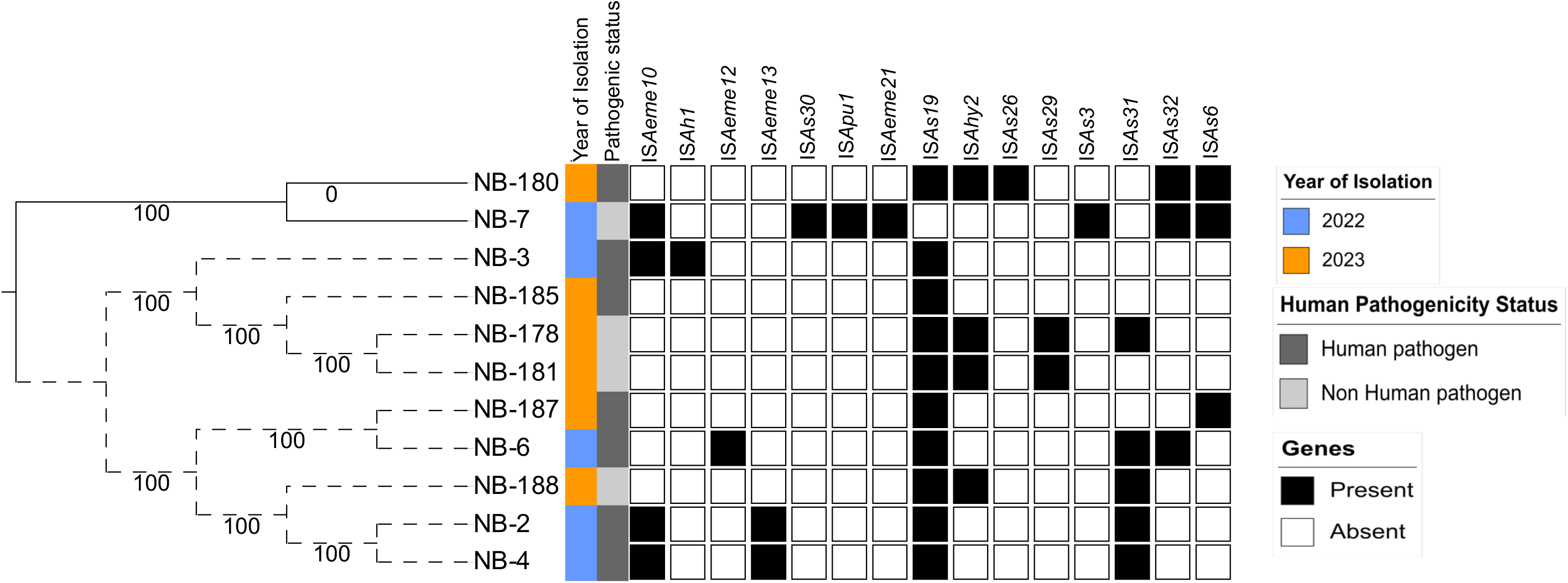
– Distribution of insertion sequence elements in *Aeromonas veronii* recovered from freshwater lake in Ontario. Core genome sequence alignment was generated from draft genomes using MAFFT v7.453, and the tree was constructed using FastTree v2.1.11 using generalized time reversible nucleotide substitution model with 1000 random resampling for node support. The dotted line depicts *A.* veronii. NB-7 (*A. allosaccharophila*) and NB-180 (*A. caviae*) were outgroups used to re-root the tree. Mobile elements were detected using Mob-suites. The scale bar at the bottom represents nucleotide substitution per site. The tree was visualized using iTOL; (https://itol.embl.de).

### Biosynthetic gene cluster profile in *A. veronii*

The 53 biosynthetic gene clusters (BGCs) identified in all sequenced *Aeromonas* species were categorized into eight BGC families using sequence similarity network analysis with BiGSCAPE (50). The ribosomally synthesized and post-translationally modified peptides (RiPPs) were the most predominant BGC class, consisting of three gene families. In contrast, the non-ribosomal polyketide synthase (NRPS) included only one gene family. The remaining gene families were classified as “others” and included homoserine lactone (n=2) and aryl polyene (n=2). The three RiPPs detected were unique and conserved within the collection but exhibited low similarity scores to previously described BGCs. For example, RiPP-1 (Figure 5a), comprising 11 open reading frames (ORFs), had a similarity score of 0.17 to angustmycin A/B/C (BGC0002621) described in *Streptomyces angustmyceticus* (Accession MZ151497.1) (51). Meanwhile, RiPP-2 and RiPP-3 (Figure 5b, Figure 5c), consisting of nine and seven ORFs respectively, had similarity scores of ≤0.08 to pseudopyronine A/B (BGC0001285) described in *Pseudomonas putida* (Accession KT373879.1) (52). Notably, RiPP-3 was also detected in *A. allosaccharophila* (NB-7), indicating that this BGC is not exclusive to *A. veronii* (Figure 5c). The identified NRPS had the highest similarity score of 0.9 to enterobactin (BGC0000343) previously described in *Pseudomonas* sp. J465 (Accession GQ370384.1) (53). This BGC was conserved in the *A. veronii* sequenced (Figure 5d). Further analysis of global *A. veronii* genomes confirmed that this BGC was conserved not only in this collection but also in all publicly available *A. veronii* genomes. A BGC encoding homoserine lactone, predominant in *A. veronii* (n=6/9), was also detected in *A. allosaccharophila*. This BGC had a low similarity score (0.14) to thioguanine (BGC0001992) in *Erwinia amylovora* CFBP1430 (Accession number: NC_013971.1) (54). Notably, a pair of *A. veronii* strains—one pathogenic (NB-6/NB-187) and one non-pathogenic to humans (NB-178/NB-181)—carried unique BGCs encoding aryl polyene (Figure 6a-b). The pathogenic pair consisted of 17 ORFs with a similarity score of 0.44 to aryl polyene (BGC0002008) described in *Xenorhabdus doucetiae* (Accession NZ_FO704550.1) (55), while the non-pathogenic pair contained 37 ORFs with a similarity score of 0.26 to bacilysin (BGC0000888) described in *Bacillus* sp. CS93 (Accession number: GQ889493.1) (56). Overall, *A. veronii* harbored putative unique BGCs that exhibited low similarity scores to previously described compounds.

**Figure 5.**
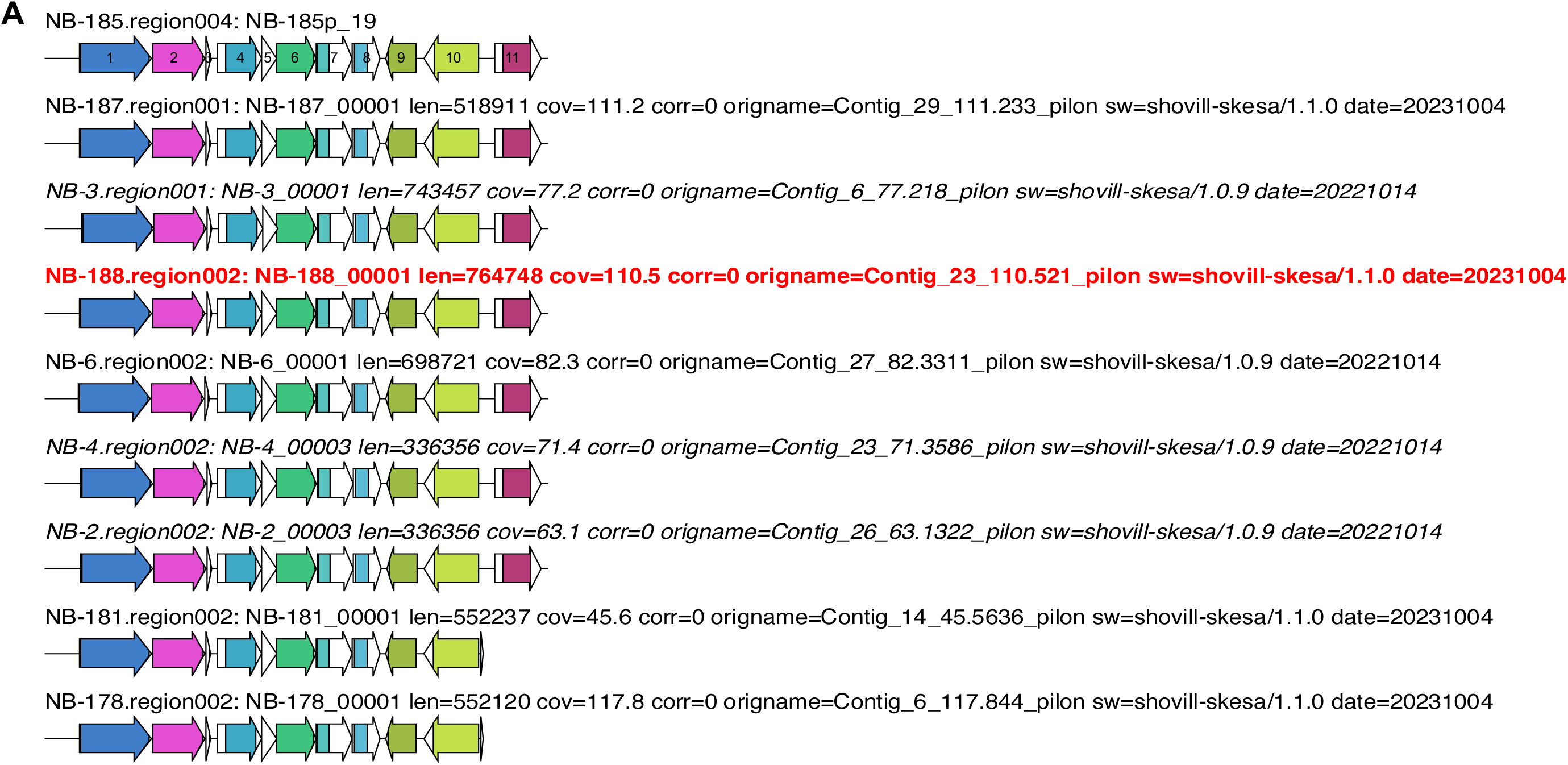
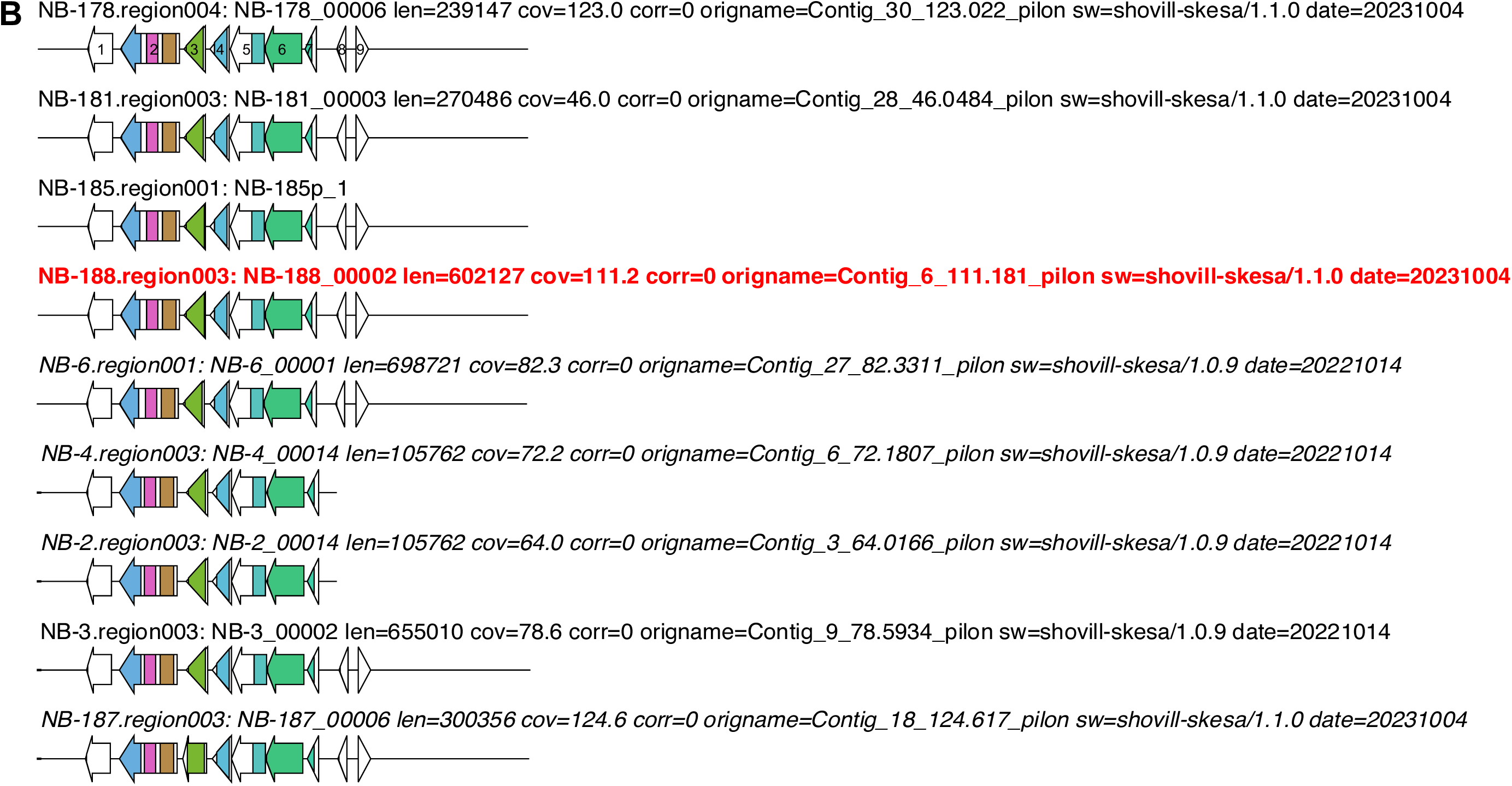
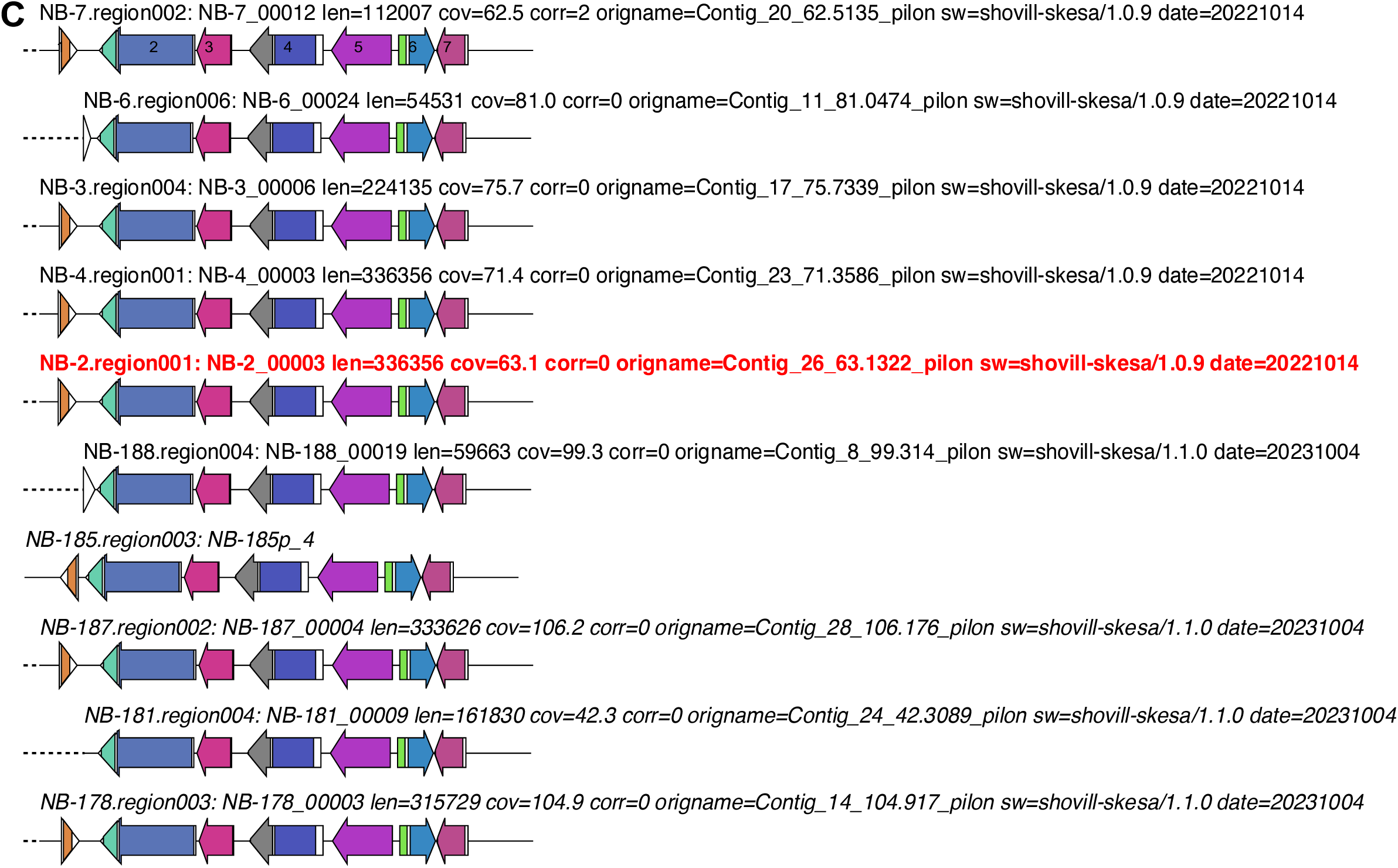
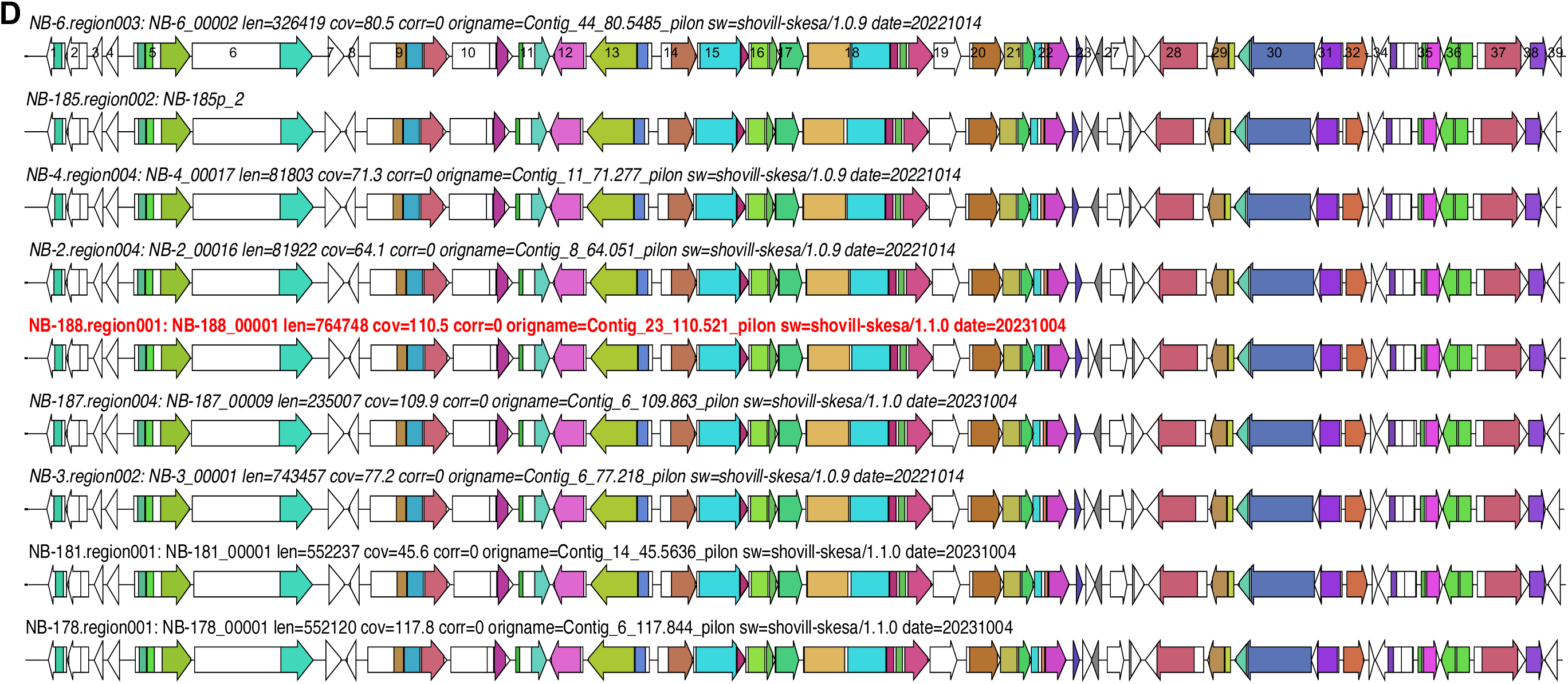
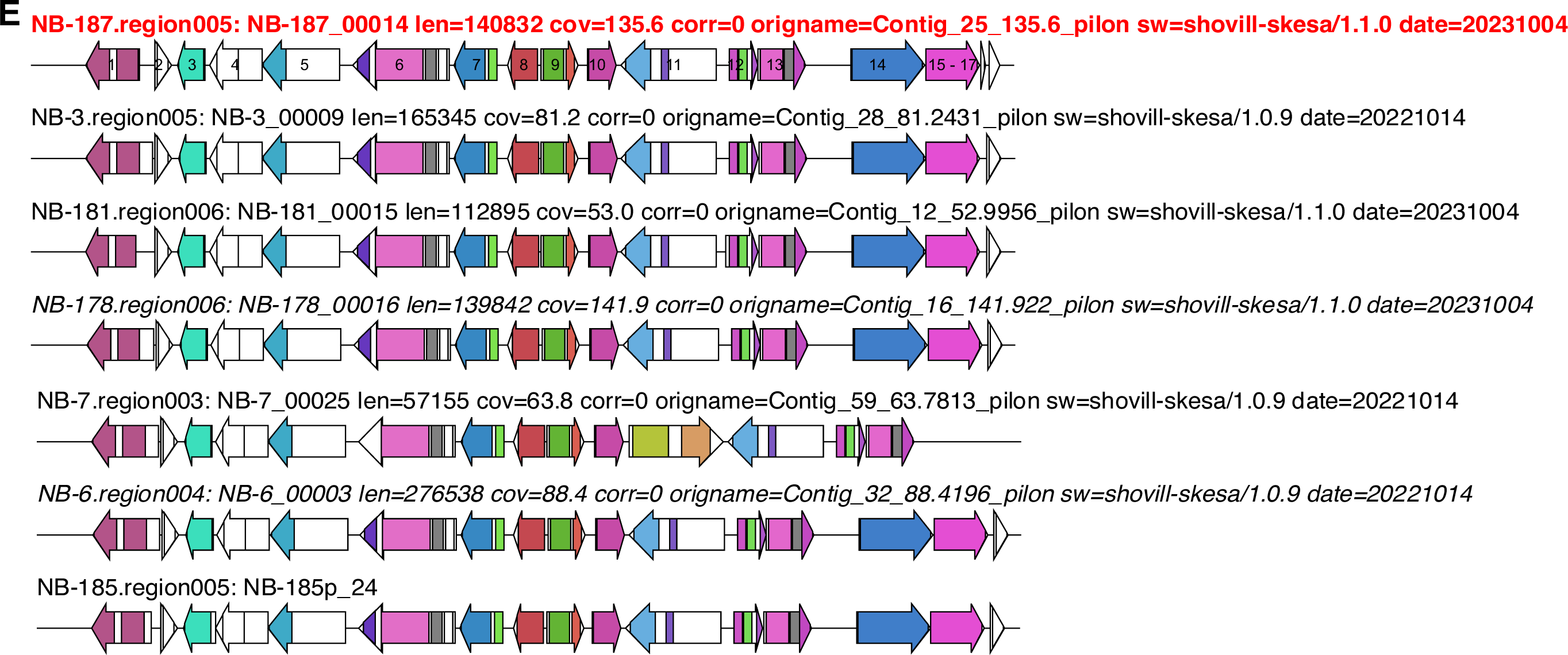
– Novel biosynthetic gene clusters identified in *Aeromonas veronii* sequenced. **A**. ribosomally synthesized and post-translationally modified peptides (RiPP) biosynthetic gene cluster (BGC) with low similarity score of 0.17 to angustmycin A/B/C in *Streptomyces angustmyceticus* (Accession MZ151497.1). **B**. RiPP gene cluster with low similarity score of 0.04 pseudopyronine A/B found in *Pseudomonas putida* (Accession KT373879.1). **C**. RiPP gene cluster detected in *A. veronii* and *A. allosaccharophila* with low similarity score of 0.08 to pseudopyronine A/B found in *Pseudomonas putida* (Accession KT373879.1). **(D).** Non-ribosomally polyketide synthase (NRPS) BGC with 0.9 similarity to enterobactin in *Pseudomonas* sp. J465 (Accession GQ370384.1). **E**. Homoserine lactone detected in *A. veronii* and in *A. allosaccharophila* with low similarity score of 0.14 to thioguanine found in *Erwinia amylovora* CFBP1430 (Accession number: NC_013971.1).

**Figure 6.**
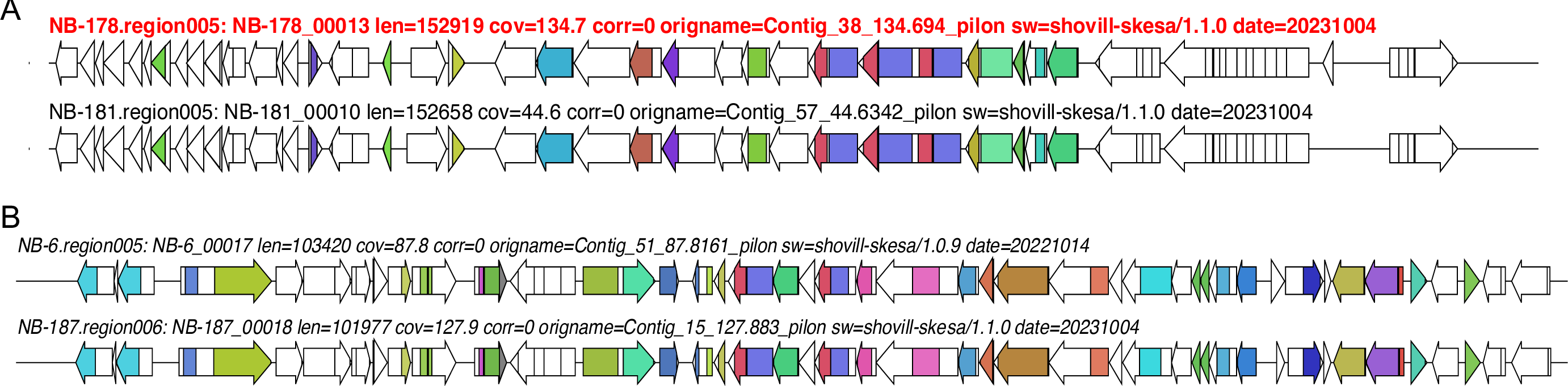
– Biosynthetic gene clusters identified in *Aeromonas veronii* predicted to be human or non-human pathogens. (A). Aryl polyene gene cluster detected in *A. veronii* isolates predicted to be non-human pathogen with similarity score of 0.26 to bacilysin found in *Bacillus* sp. CS93 (Accession number: GQ889493.1). (B). Aryl polyene gene cluster detected in *A. veronii* isolates predicted to be human pathogen with similarity score of 0.44 to aryl polyene found in *Xenorhabdus doucetiae* (Accession number: NZ_FO704550.1).

## DISCUSSION

Aquatic ecosystems are continually impacted by anthropogenic activities, making the microbial quality and safety of these water bodies, especially those used for recreational activities, paramount for public health (1). In this study, we assessed the presence of *A. veronii* in a recreational lake and determined the extensive genomic features of the isolates regarding their population structure and the genomic characterization of stress response genes, mobile genetic elements, and other gene content such as biosynthetic gene clusters that confer uniqueness to different *A. veronii* strains. We inferred the global population structure of *A. veronii* by assessing the genetic relatedness of the isolates sequenced with previously sequenced strains in public databases.

In the past decade, WGS has become the gold standard method for species identification, complementing existing biochemical-based methods (57). In this study, the WGS-based approach identified isolates as *A. veronii, A. caviae*, and *A. allosaccharophila*, whereas the biochemical identification system misidentified all isolates as either *A. sobria* or *A. hydrophila*. Misidentification of species of environmental bacteria by biochemical approaches is not uncommon (57). Studies comparing biochemical-based bacterial species identification systems to WGS have shown that species misidentification can vary by species and is common in specific bacteria, including *Pseudomonas fluorescens*, *Pseudomonas putida* (57), and *Enterococcus faecalis* (58).

*A. veronii* strains sequenced exhibited different pathogenic potentials, with the majority (67%, n=6/9) predicted to be pathogenic to humans and possessing a similar virulence determinant profile, including T3SS. T3SSs are crucial virulence mechanisms that allow bacteria to inject effector proteins directly into the host cell cytoplasm. The activity of T3SSs closely correlates with infection progression and outcome in various infection models, and its presence is considered a general indicator of virulence in *A. veronii* (59–61). The detection of human pathogenic *A. veronii* in this study, along with other clinically relevant pathogens such as *Bacillus anthracis* (21) and *Vibrio cholerae* (22) in this recreational lake from previous studies, emphasizes the crucial role aquatic ecosystems play in disseminating pathogens. The recreational use of this water could pose a continuous risk to public health, serving as a reservoir and facilitating the transmission of waterborne diseases. This also underscores the significance of monitoring aquatic environments as reservoirs for pathogenic bacteria.

There was high genetic diversity among the nine *A. veronii* isolates sequenced, including the identification of novel sequence types and alleles. While some strains were indistinguishable by SNPs, others were genetically distant. This finding could imply that different *A. veronii* strains may have been introduced into the lake multiple times from various sources such as resident freshwater fish, domestic animals, and environmental samples (5). The integration of genomic data from the *A. veronii* isolates sequenced with global strains revealed that isolates from single sites formed smaller groups within the phylogeny. Interestingly, one isolate from this study (NB-188) was nested with an isolate recovered from an aquatic ecosystem in France. A previous study assessing the core genome-based phylogenetic analysis of *A. veronii* genomes deposited in NCBI from 18 countries revealed high genetic diversity (5). The admixture of *A. veronii* strains from different sources was observed, suggesting a lack of source- and timepoint-based clustering in the *A. veronii* population. However, strains from a single site tend to form small groups within the phylogenetic clusters. These observations concur with our findings. The genetic diversity observed in *A. veronii* reinforces the importance of continuous genomic surveillance to monitor the emergence and spread of virulent and/or resistant strains.

The AMR determinant profile observed in the isolates sequenced in this study was comparable and included only chromosome-borne genes encoding resistance to beta-lactams. The prevalence of β-lactam resistance genes, including those conferring resistance to carbapenems, is a known phenomenon in the *A. veronii* population (10). While these genes were chromosomal with no close proximity to mobile genetic elements, their spread to other strains or bacterial species is not entirely unlikely as bacterial cell lysis could release DNA into the environment where it could be taken up by other strains or bacterial species through the process of transformation. Indeed, natural transformation has been described as a common mechanism of horizontal gene transfer among *Aeromonas* species, including *A. veronii. Aeromonas* species are capable of competence and transformation (62). In addition, *A. veronii* is known to easily acquire and exchange AMR genes (7, 20, 63). Although there was a low occurrence of AMR in *A. veronii* in this study, Lake Wilcox is a potential reservoir for AMR genes encoding resistance to multiple antibiotics as evidenced by results from previous studies on the lake where other bacterial species isolated from the lake carried multiple AMR genes (21, 22).

The mobilome is known to facilitate gene gain and loss, a phenomenon that plays a crucial role in bacterial evolution and ecological adaptation and a probable change in bacterial fitness (64, 65). This change can contribute to the emergence of divergent bacterial populations with unique features, including higher pathogenic potential (64, 66, 67). In this study, no plasmid was detected in the *A. veronii* sequenced, but other mobile genetic elements (MGEs) including prophages and insertion sequences were identified. The majority of the intact prophages were identified as P2-like phages (Peduoviridae) (46), and a few of them were predicted to have multiple host bacterial species. This observation is interesting and could suggest a broad host range of these phages, which could have applications in biocontrol (68–70), but further studies on the host range of these phages would be needed to ascertain this. Another factor that contributes to the rapid evolution and ecological adaptation and that could influence the pathogenicity of bacterial species is BGCs that encode the production of various secondary metabolites (52, 71, 72). This phenomenon is seldom studied in *A. veronii*.

In this study, we found a high abundance of novel BGCs and identified unique NRPS and RiPP that were conserved in *A. veronii.* Notably, NRPS with high similarity (0.9) to enterobactin found in *Pseudomonas* sp. J465 (53), that mediates high affinity for iron acquisition in stringent conditions (73, 74). Angustmycin A/B/C (51) and pseudopyronine A/B (52) homologs were found to be conserved in *A. veronii.* These RiPP products encode antimicrobial properties and contribute to the survival of their producers in their ecological niche (51, 75). These conserved clusters could be promising genomic markers for typing *A. veronii*. Of note, a bacilysin homolog gene (56) was detected in a pair of non-human pathogenic strains. Bacilysin is an antimicrobial dipeptide produced by *Bacillus* species that exhibits antagonistic activity against both Gram-negative and Gram-positive bacteria (56, 76, 77). Further studies would be required to decipher the antimicrobial activity of the bacilysin homolog identified in this study against human pathogenic strains of *A. veronii* and other pathogens, as well as their mechanism of actions.

## CONCLUSION

The study presents a genomic analysis of *A. veronii* strains isolated from a freshwater lake and defined the population structure and characterized the genetic factors associated with stress and ecological adaptation. A significant finding is the pathogenic potential of *A. veronii* to humans that underscores the public health implications, especially considering the recreational use of the lake. Among the MGEs identified that could contribute to the genetic diversity, adaptability, and the pathogenicity to human, as well as the of *A. veronii* as a reservoir for AMR genes, the BGCs identified presents opportunities for the discovery of novel bioactive compounds. Overall, this study not only contributes to our understanding of the genetic diversity and ecological dynamics of *A. veronii* but also highlights the potential public health risks and AMR reservoir role of this bacterium. It underscores the need for continuous surveillance for pathogens in aquatic ecosystems.

## Funding

We acknowledge support from the Canada First Research Excellence fund in support of this project.

## Authors’ contribution

OL, NB, VP, RA, MS, YC, MP conducted the sampling, isolation, and whole-genome sequencing; OL performed the bioinformatics analysis and wrote the original draft of the manuscript. LG conceived the project and provided funding and resources. OL, VP and LG supervised the study. All authors read and approved the final manuscript.

## Legends

**Table 1 -** Summary of sequence metrics of *Aeromonas* isolates recovered from a freshwater lake.

**Table 2** – Features of intact phages detected in *Aeromonas veronii* sequenced in the study.

## Supplementary Data

**Table S1** - Summary of the novel alleles and sequence types identified in *Aeromonas* isolates sequenced in this study.

**Table S2** - Distance matrix between *Aeromonas veronii* isolates sequenced in this study.

**Table S3** - List of accession number and associated metadata of publicly available *Aeromonas veronii* genomes retrieved from the GenBank NCBI database.

## Notes

### Competing Interest Statement

The authors have declared no competing interest.

